# Constrained sampling from deep generative image models reveals mechanisms of human target detection

**DOI:** 10.1101/578633

**Authors:** Ingo Fruend

## Abstract

The first steps of visual processing are often described as a bank of oriented filters followed by divisive normalization. This approach has been tremendously successful at predicting contrast thresholds in simple visual displays. However, it is unclear to what extent this kind of architecture also supports processing in more complex visual tasks performed in naturally looking images.

We used a deep generative image model to embed arc segments with different curvatures in naturalistic images. These images contain the target as part of the image scene, resulting in considerable appearance variation of target as well as background. Three observers localized arc targets in these images, achieving an accuracy of 74.7% correct responses on average. Data were fit by several biologically inspired models, 4 standard deep convolutional neural networks (CNN) from the computer vision literature, and by a 5-layer CNN specifically trained for this task. Four models were particularly good at predicting observer responses, (i) a bank of oriented filters, similar to complex cells in primate area V1, (ii) a bank of oriented filters followed by tuned gain control, incorporating knowledge about cortical surround interactions, (iii) a bank of oriented filters followed by local normalization, (iv) the 5-layer specifically trained CNN. A control experiment with optimized stimuli based on these four models showed that the observers’ data were best explained by model (ii) with tuned gain control.

These data suggest that standard models of early vision provide good descriptions of performance in much more complex tasks than what they were designed for, while general purpose non-linear models such as convolutional neural networks do not.

## Introduction

Processing in higher levels of the visual system becomes successively invariant to the specific details of an image. For example, orientation tuning in primary visual cortex seems to be largely independent of stimulus contrast (Finn et al., 2007; Nowak & Barone, 2009; Skottun et al., 1987; Bowne, 1990, at least for large stimuli Mareschal & Shapley, 2004), shape processing is invariant to rotations (Blais et al., 2009), and specifically object recognition seems invariant to many different image transformations, including contrast, rotation, scaling, and illumination (Gauthier & Tarr, 2016; Pinto et al., 2007; DiCarlo et al., 2012). The visual system derives invariance with seemingly no effort even in natural scenes, with lots of clutter and possible occlusions (Balboa et al., 2001; DiMattina et al., 2012) and despite often dramatic variations in appearance (Elder & Zucker, 1998).

Over the years, a standard model for the early stages of visual processing has emerged, that consists of a bank of linear filters tuned to image properties like spatial frequency (Campbell & Robson, 1968; Stromeyer & Julesz, 1972) and orientation (Wilson et al., 1983) followed by divisive gain control (Legge & Foley, 1980; Foley, 1994; Heeger, 1992; Albrecht & Geisler, 1991; Mante et al., 2005; Schütt & Wichmann, 2017; Neri, 2015; Foley, 2019). This theory can successfully explain the independence of orientation tuning and contrast (Finn et al., 2007) and it generalizes over a wide range of different stimulus conditions (Neri, 2015). So far, however, it is unclear if it can also account for higher levels of visual processing and for more complex invariances.

One challenge here is the difficulty of generating and manipulating stimuli with naturalistic variability in appearance. Modern generative image models based on artificial neural networks, such as Generative Adversarial Networks (GANs, Goodfellow et al., 2014), can generate impressively realistic images (see examples in Brock et al., 2018; Miyato et al., 2018) and human visual performance is highly sensitive to image manipulations guided by these models (Fruend & Stalker, 2018).

Zhu et al. (2016) propose a method to edit images directly within the manifold of natural images, by applying user defined constraints on images sampled from a GAN. Here, we use their method to embed edges in artificial, yet natural looking images and we ask human observers to detect these edges. It is well known that edges in natural images have a considerable degree of appearance variation (Elder & Zucker, 1998) and the embedding method by Zhu et al. (2016) is expected to (at least partly) reproduce this variability.

Artificial neural networks have recently attracted interest in the vision science community because they appear to successfully perform object recognition in photographic images (Krizhevsky et al., 2012; Simonyan & Zisserman, 2015; He et al., 2015). Furthermore, artificial neural networks have been argued to make similar errors as humans in tasks that involve intermediate visual representations (Kubilius et al., 2016) and to develop internal representations that seem to resemble those within the primate ventral stream (Khaligh-Razavi & Kriegeskorte, 2014; Yamins et al., 2014). We therefore compare our observer’s detection performance to a number of variations of the standard model as well as several artificial neural network models.

## Materials and methods

### GAN training

We trained a Wasserstein-GAN (Arjovsky et al., 2017) on the 60 000 32*×*32 images contained in the CIFAR10 dataset (Krizhevsky, 2009) using gradient penalty as proposed by Gulrajani et al. (2017). In short, a GAN consists of a generator network *G* that maps a latent vector ***z*** to image space and a discriminator network *D* that takes an image as input and predicts whether that image is a real image from the training dataset or an image that was generated by mapping a latent vector through the generator network (see Figure 1). The generator network and the discriminator network were trained in alternation using stochastic gradient descent. Specifically, training alternated between 5 updates of the discriminator network and one update of the generator network. Updates of the discriminator network were chosen to minimize the loss

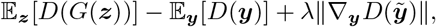

and updates of the generator were chosen to maximize this loss. Here, the first term quantifies the false alarm rate of the discriminator (i.e. the likelihood that the discriminator *D* classifies a fake image *G*(***z***) as real), the second term quantifies the hit rate of the discriminator and the third term is a penalty term to encourage the discriminator to be 1-Lipschitz (a stronger form of continuity). In accordance with Gulrajani et al. (2017), we set *λ* = 10 for discriminator updates and *λ* = 0 for generator updates. In this equation, 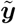 denotes a random location between ***ŷ*** = *G*(***z***) and the training image ***y***. Networks with different numbers of hidden states (parameter *N* in Figure 1) were trained for 200 000 update cycles using an ADAM optimizer (Kingma & Ba, 2015) with learning rate 10^*−*4^ and *β*_0_ = 0, *β*_1_ = 0.9. Specifically, we trained networks with *N* = 40, 50, 60, 64, 70, 80, 90, 128 (see Figure 1). Wasserstein-1 error (Arjovsky et al., 2017) on a validation set (the CIFAR10 test dataset) was lowest with *N* = 90 in agreement with visual inspection of sample quality, so we chose a network with *N* = 90 for all remaining analyses. In Appendix “Naturalism of CIFAR10 images” we provide evidence that the CIFAR10 dataset contains image features that a naïve observer would likely use during natural vision.

**Figure 1:**
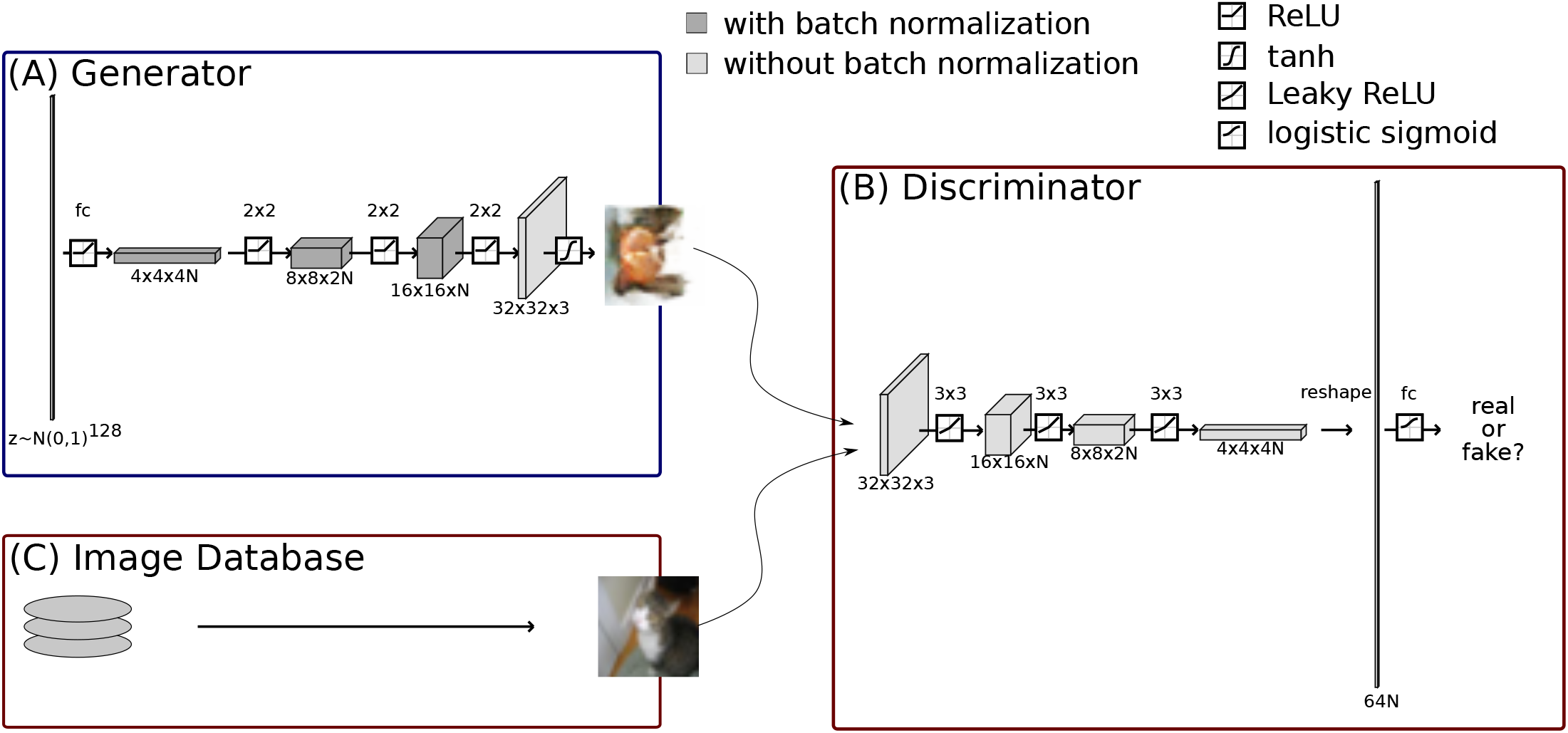
Architecture of the generative adversarial network. (A) Architecture of the generator network. Information flow is from left to right, each array corresponds to one transformation of the input. The square insets on the arrows indicate the respective non-linearity (see upper right for legend), the labels above the arrows indicate the kind of affine transformation that was applied (fc: fully connected, i.e. unconstrained affine transformation, 2 *×* 2 transpose convolution, i.e. upsampling before convolution to increase spatial resolution and image size). Blocks indicate hidden unit activations. For darker blocks, batch-normalization (Ioffe & Szegedy, 2015) was applied, for lighter blocks, no batch-normalization was applied. “ReLU” refers to rectified linear unit “ReLU(*x*) = max(0, *x*)” (Glorot et al., 2011). The generator network maps a sample *z* from an isotropic 128 dimensional Gaussian to a 32 *×* 32 pixel color image. (B) Architecture of the discriminator network. Same conventions as in (A), but 3 *×* 3 is for regular convolution with a stride of 2 and kernel size of 3. See He et al. (2015) for definition of “leaky ReLU”. The discriminator network receives as input either an image *ŷ* generated by the generator network or a real training image *y*, from the image database (C) and it decides if the input image is real or not.

### Observers

Three observers gave their informed consent to participate in the study. All observers were naïve to the purpose of this study. Observers o1 and o2 had some experience with psychophysical experimentation, while o3 did not. All observers had normal or corrected to normal vision were between 23 and 26 years old and received no monetary compensation for participation. The study procedures were approved by the Ethics Committee of York University, Toronto, ON and adhered to the principles outlined in the original Declaration of Helsinki.

### Stimuli

Stimuli were 64*×*64 pixel grayscale images presented at 8bit resolution on a Sony Triniton Multiscan G520 CRT monitor in a dimly illuminated room. The stimulus images were initially constructed at 32*×*32 pixel resolution and were subsequently upsampled for display using bilinear interpolation. The monitor was carefully linearized using a Minolta LS-100 photometer. Maximum stimulus luminance was 106.9 cd/m^2^, minimum stimulus luminance was 1.39 cd/m^2^. The background was set to medium gray (54.1 cd/m^2^). At the viewing distance of 56 cm, the stimulus square subtended approximately 1 degree visual angle. One pixel subtended approximately 0.031 degree of visual angle. Two different types of stimuli were used, arc segments embedded in naturalistic images and model optimized stimuli to target different performance levels.

#### Embedding arc segments in naturalistic images

Stimuli in the main experiment consisted of arc segments that were embedded into naturalistic images by conditionally sampling from a GAN. In the 32*×*32 pixel image array, each arc segment connected one of four pairs of points. These were (13, 20) *−* (20, 20) (top location), (13, 20) *−* (13, 13) (left location), (13, 13) *−* (20, 13) (bottom location), or (20, 13) *−* (20, 20) (right location), where coordinates are in pixels from the bottom left corner of the image (see Figure 3). Furthermore, arc segments had one of eight curvatures 0, *±*0.096, *±*0.177, *±*0.231, 0.25, corresponding to orientations of *±nπ*/8, *n* = 0, …, 7 at their endpoints. These arcs were then rendered to a 32*×*32 pixel image using a background intensity of 0 and a foreground intensity of 1. The differentiable histogram of oriented gradients (HOG) by Zhu et al, (Zhu et al., 2016) was evaluated on each one of them.

In order to sample an image from the GAN that contained the respective arc segment, we follow Zhu et al. (2016) and minimized the objective function

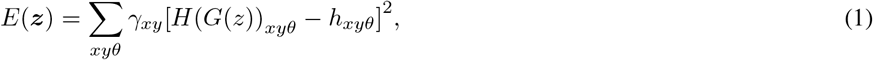

where the sum goes over all pixel locations *x, y* and 8 different equally spaced orientations *θ*, *H*: ℝ^32*×*32^ → ℝ^32*×*32*×*8^ is the differentiable HOG operator, *G*: ℝ^128^ → ℝ^32*×*32^ is the generator network of the GAN and *h* = (*h*_*xyθ*_) ∈ ℝ^32*×*32*×*8^ is the differentiable HOG operator evaluated on the target arc image. Thus, the minimized error function was simply the squared difference between local histograms of oriented gradients in the generated image and in the target binary arc image. To make sure that only errors in the vicinity of the embedded arc could drive image generation, we weighted the error terms by weights *γ*_*xy*_ that were calculated by convolving the binary arc image with a kernel of the form

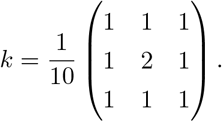

This way error terms arising from pixels that were more than 2 pixels away from the embedded arc received a weight of zero and did not drive the search for an appropriate latent vector ***z***_arc_. To find a latent vector ***z***_arc_ that corresponded to an image that contained the respective arc segment, we started with a random latent vector ***z***_0_ drawn from an isotropic Gaussian distribution and then used gradient descent on *E*(***z***) as defined in Equation (1). While Zhu et al. (2016) aimed for real time performance and therefore trained a neural network to optimize equation (1) in one shot, we did not require real time performance and decided to simply minimize equation (1) using gradient descent. In Appendix “Validation of embedded arc stimuli as “natural””, we show that the non-target areas of the constructed stimuli were statistically very similar to natural images from the CIFAR10 database.

We constructed a total of 134 400 stimuli (4 locations, 8 curvatures, 4200 exemplars, see Figure 2 for examples).

**Figure 2:**
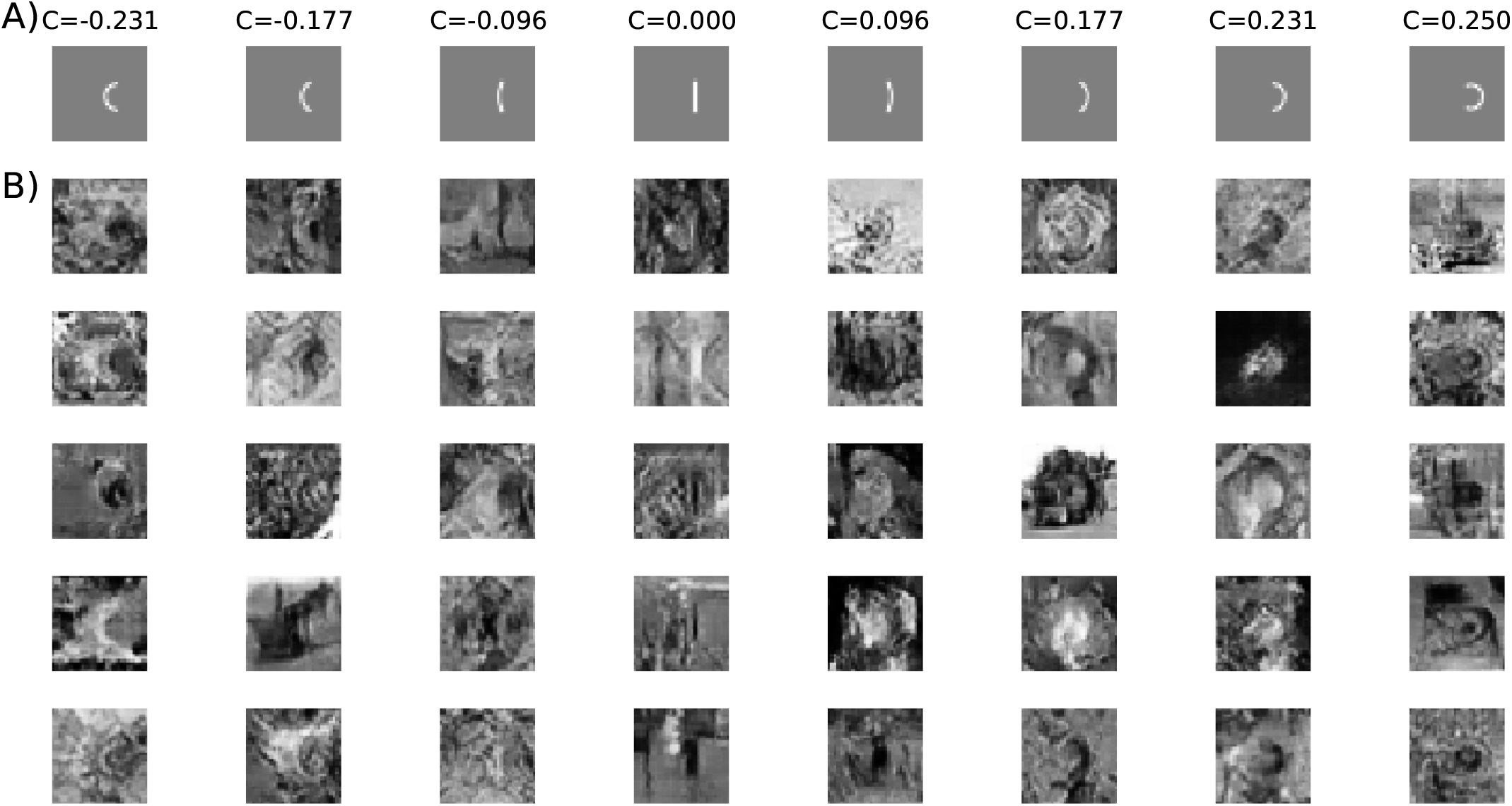
Embedding an arc segment in a natural image. (A) Arc stimuli embedded in the images. Only the “right” location is shown. The corresponding curvature values are shown above the images. (B) Natural images with embedded arc segments. All stimuli in one column correspond to the same arc segment.

#### Optimizing stimuli to target different performance levels

In a second experiment, we tested observers’ performance on stimuli that were optimized with respect to a candidate model (see Section “Model fitting” and Appendix “Details of model architectures” for details of these models). Specifically, let 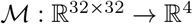 be a candidate model that maps a 32*×*32 pixel image *I* to a vector of probabilities 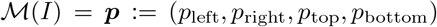. To construct a stimulus for which the model predicts a correct left-response with probability *q*, we minimized the error function

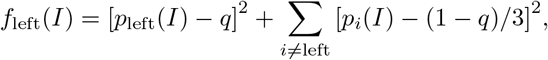

and similar for right, top, and bottom responses. For each location and for target performances of *q* = 25%, 50%, 75%, 95%, we constructed 40 stimuli using gradient descent on the image pixels.

Optimized stimuli were only computed for the four best models (see Section “Prediction performance can not discriminate between different models”).

### Procedure

We performed two experiments that were the same in everything but the stimuli presented (see Section “Stimuli”). All other procedures were the same between both experiments.

Figure 3 shows the layout of a single trial. Each trial started with an 800ms fixation interval. During the fixation interval, a marker consisting of four dots (size 1 pixel) forming the corners of a 10*×*10 pixel (0.31*×*0.31 degree visual angle) square was visible on the screen and observers were instructed to fixate at approximately the center of the square. After the fixation interval, the target stimulus appeared on the screen for 100ms. On all experimental trials, the target stimulus was followed by a second interval showing only the fixation marker and observers had one second to indicate where they had seen the arc segment by pressing the corresponding arrow key on a computer keyboard; the up-key if the arc segment was between the top two dots, the down-key if the arc segment was between the bottom two dots, the left-key if the arc segment was between the left two dots, and the right-key if the arc segment was between the right two dots. After the response, observers always received feedback: After a correct response, the fixation marker jumped up for a duration of 9 frames (150ms), after an incorrect response, the fixation marker jiggled randomly for a duration of 9 frames (150ms). If observers did not respond within one second, the message “no response” was flashed on the screen and the trial was marked as *invalid response* and was discarded from further analysis. This affected between 57 and 61 trials per observer (approximately 1% of all trials).

**Figure 3:**
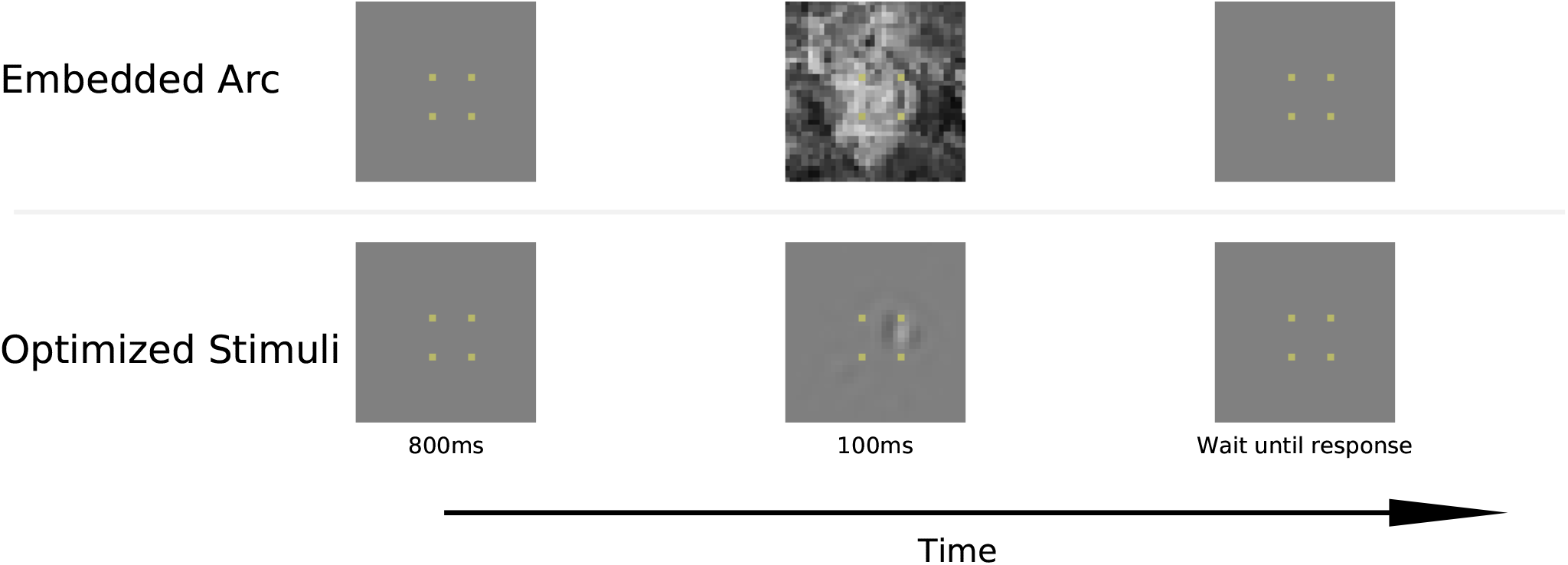
Example experimental display. Top row: Trial sequence in the embedded arc experiment. Bottom row: Trial sequence in the optimized stimuli experiment with an example stimulus optimized for the Feature Normalization model for observer o1. Time increases from left to right. In both cases, the correct target response would be “right”. The dots mark the end points of the possible arc segments and were also present during the experiment.

At the beginning of the experiment, each observer performed 20 trials of training, in which they saw the isolated arc segments and had an opportunity to familiarize themselves with the response format. After that, observers performed two sessions of 400 trials each for training with the embedded arc stimuli. In the first training session, observers performed the task on unmasked stimuli. In the second training session, observers performed the task on masked stimuli; after 100ms, a second, unconditioned sample from the GAN appeared on the screen and remained visible for up to one second or until the observer made a response.

After the training sessions, each observer performed 8 sessions with 400 trials each with embedded arc segment stimuli, without masking. During these 8 sessions, performance did not change considerably as confirmed by plotting a rolling 20 trial response accuracy. We then performed two additional control experiments. In the first control experiment, the stimuli from one of the sessions from the main experiment were shown a second time. This allowed us to determine how consistent observers responded with their own responses (Neri & Levi, 2006). In the second control experiment, we used stimuli that were optimized to target different performance levels for model observers. This second control experiment used the stimuli described in Section “Optimizing stimuli to target different performance levels”. In the second control experiment, observers saw 10 trials for each combination of arc-location, target performance, and model.

### Model fitting

We evaluated eleven different models for their ability to predict human responses on a trial-by-trial level. The first 6 models were based on a bank of orientated filters with parameters that mimicked spatial tuning properties of neurons in primary visual cortex (Ringach et al., 2002). These models differed between each others in two aspects. Firstly, models were either based on the filter outputs directly, or they were based on the energy of the filter outputs (Morrone & Burr, 1988; Adelson & Bergen, 1985) to mimic complex cell responses in primary visual cortex. Secondly, models were either directly using these filter outputs, or they applied spatially local or orientation tuned gain control to the filter outputs. We also used a number of deep convolutional neural networks (LeCun et al., 2015), allowing for highly non-linear features. Specifically, we looked at two classes of neural networks. The first class were standard neural networks with published architectures for which versions pretrained on the high resolution ImageNet database are publicly available. For these models, we kept the initial feature layers intact and re-trained a final linear decoding layer. Secondly, we specifically optimized a neural network model to predict human responses for each one of the observers. All models derive a set of (linear or non-linear) features and human responses were predicted by linearly decoding these features. Details of the models can be found in Appendix “Details of model architectures”.

Before fitting the models to the observers’ data, we split the data into a training set (80% of all trials, between 3200 and 3500 trials), a validation set (10% of all trials, between 400 and 440 trials), and a test set (10% of all trials, between 400 and 440 trials).

The biologically inspired models are ultimately logistic regression models with elaborate input features. As such, they could easily be fit using deterministic gradient based methods such as gradient descent or Fisher scoring iterations. This is however not the case for the deep convolutional neural network models. Such models are typically trained using stochastic gradient descent; the training data is split into a number of smaller “mini-batches” and gradient descent is performed using gradients of the likelihood of these small subsets of the data. This allows stochastic gradient descent to make more efficient use of the compute power and in addition, the stochasticity arising from taking the gradient on random subsets of the data potentially allows the model to escape from local minima (Murphy, 2012, p. 267). To ensure that differences between models did not result from the training method used, we decided to train all models using stochastic gradient descent on the negative log-likelihood of the training data. We used a batch size of 16 trials and a learning rate of 0.001 (except for the Energy model, for which a learning rate of 0.1 was used). Models were trained for up to 200 epochs, where one epoch consisted of one pass over all training data. We used early stopping to regularize the models (e.g. Goodfellow et al., 2016, chapter 7.8): After every epoch, we evaluated the negative log-likelihood of the validation data set. If this validation likelihood did not improve over 10 successive epochs, model training was interrupted early.

## Results

### Performance varies weakly with arc curvature

Observers’ performance did not vary much with the curvature of the embedded arc. As shown in Figure 4, all observers performed at a level of approximately 75% correct (o1: 80.1 *±* 0.67%, o2: 74.6 *±* 0.73%, o3: 69.5 *±* 0.81%). For two out of three observers, more acute arcs were slightly easier to detect than straight arcs (o1: correlation between performance and squared curvature *r* = 0.65, *p <* 0.08 Wald test, o2: *r* = 0.72, *p <* 0.05, o3: *r* = 0.90, *p <* 0.005). However, these variations only covered a relatively small performance range.

**Figure 4:**
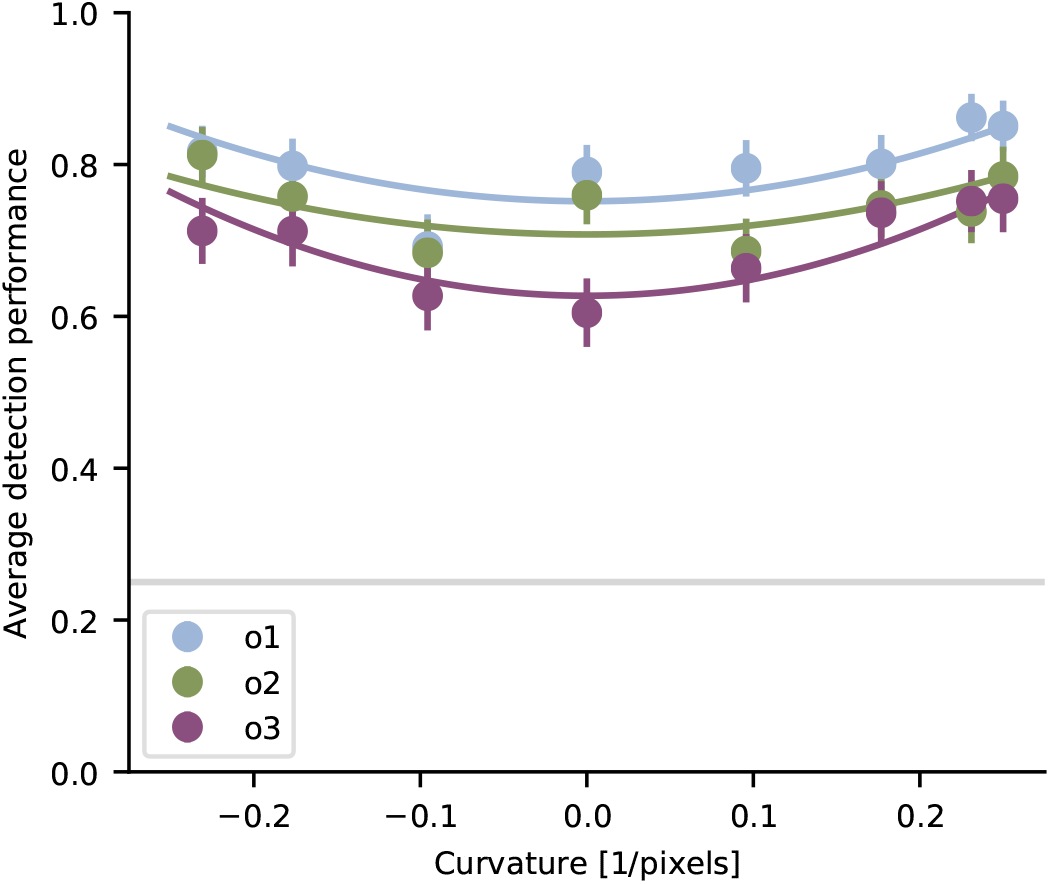
Performance for different curvatures. Mean fraction of correct responses is shown for different observers (color coded). Solid parabolas are least squares fits of the model *p*(correct) ≈ *a* + *bC*^2^. Error bars indicate 95%-confidence intervals determined from 1000 bootstrap samples. The horizontal gray line marks chance performance.

### Prediction performance can not discriminate between different models

We compared a number of different models for the trial-by-trial behaviour of observers in the arc detection task. These models consisted of a logistic regression applied to abstractions of features that are known to be computed at different levels by the visual system. As a first step of evaluating these models, we assessed how well they predicted the trial-by-trial responses of individual observers on a held out set of previously unseen trials from the same experiment.

For the first model, the features consisted of a bank of linear filters with tuning parameters that resembled those of simple cells in macaque area V1 (see Section “Model fitting” and Appendix “Details of model architectures”). These features achieved an average prediction accuracy of 51.2%, which was significantly above chance (*p <* 10^*−*10^, binomial test, see Figure 5A). However, the prediction accuracy of this linear model was still much below the internal consistency of the observers as determined by running one of the experimental sessions a second time.

**Figure 5:**
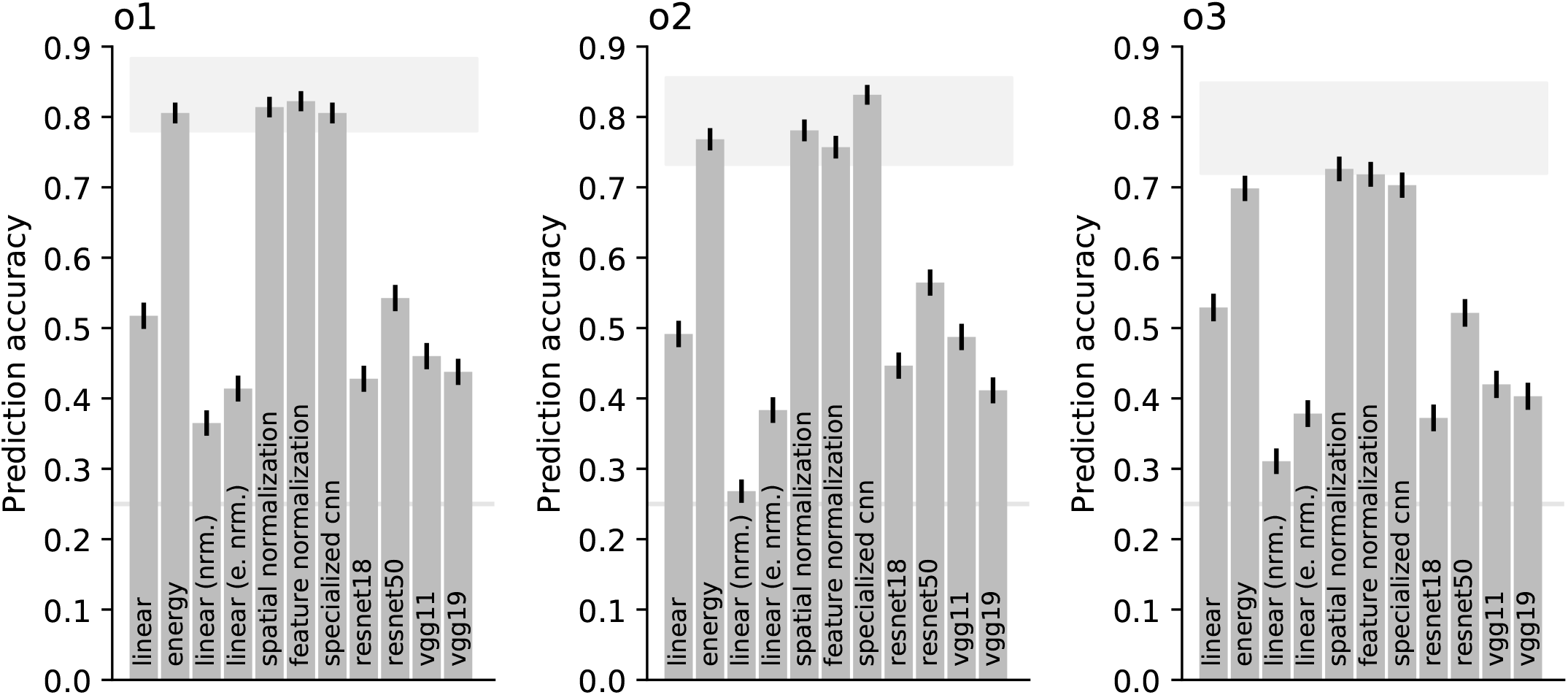
Predictive performance of evaluated models. Prediction accuracy for different models on a held out test-set of trials. Error bars indicate standard errors on the test-set. Models on the X-axis correspond to the features used by the model. The light gray line indicates chance performance, the light gray area at the top marks the range for the best possible model performance derived by Neri & Levi (2006).

The second model replaced the linear filters by energy detectors with the same tuning parameters, resembling complex cells in area V1. This model—which we will in the following refer to as the “Energy” model—achieved an average prediction accuracy of 75.7%, similar to the observers’ internal consistency.

We furthermore explored a number of models that applied different kinds of normalization to either the linear filters or the energy filters. In general, these models used either a linear filter bank or a bank of energy detectors and normalized their responses by responses that were either pooled locally in space or in feature space. These models generally replicated the observations made with the first two models: Normalized energy features achieved performances that resembled the internal consistency of the observers, while normalized linear features did not. Furthermore, we found that models based on local energy with normalization by locally pooled energy (the “spatial normalization” model in Figure 5) or by energy pooled in feature space (the “feature normalization” model in Figure 5) performed as good or slightly better than the simple energy model (o2: specialized cnn - energy = 6.3%, *p <* 1*/*2000, permutation test with 2000 samples^1^, o3: spatial normalization - energy = 2.7%, *p <* 1*/*2000, all other tests n.s.). Yet, models based on normalized linear features performed worse than the model based on unnormalized linear features (accuracy differences *>* 10%, *p <* 1*/*2000 permutation test with 2000 samples).

We also explored a number of models with hierarchically constructed, highly non-linear features. These models were implemented as multilayer convolutional neural networks. Four such models were based on successful versions of models from the computer vision literature (see Appendix “Details of model architectures” for details). The features used by these models were optimized to aid object recognition on a large database of natural images. These models are labelled as resnet18 and resnet50 (He et al., 2015) and as vgg11 and vgg19 (Simonyan & Zisserman, 2015) in Figure 5. Notably, these models are not able to predict human performance very well and achieve prediction accuracies between 41 and 55% on average. This is not overly surprising given that the task crucially relied on a model’s ability to predict the *location* of the target. In contrast, these models have been optimized to predict the *identity* of a target and they seem to do so by largely ignoring precise spatial structure (Brendel & Bethge, 2019). However, when specifically training a convolutional neural network to predict human trial-by-trial performance in our task, such a model was indeed able to achieve performance levels that were similar to the models based on energy detectors (specialized cnn model in Figure 5), so cnns are in principle able to do this task.

Neri & Levi (2006) found that the best trial-by-trial prediction performance that any model could achieve depends on assumptions about an observer’s internal noise. However, they show that the best trial-by-trial prediction performance is always between the observer’s internal consistency and an upper bound as marked by the light gray areas in Figure 5. Four models were within (or in case of o3 very close to) this range; the Energy model, the Spatial Normalization model, the Feature Normalization model and finally the specifically trained convolutional neural network model.

### Targeted stimuli reveal advantage of standard model

In order to discriminate between the different candidate models, we constructed artificial stimuli that directly targeted specific performance levels for each of the top models from Figure 5 (Energy, Spatial Normalization, Feature Normalization, CNN). Examples of these stimuli for observer o1 are shown in Figure 6 (stimuli for other observers were similar).

**Figure 6:**
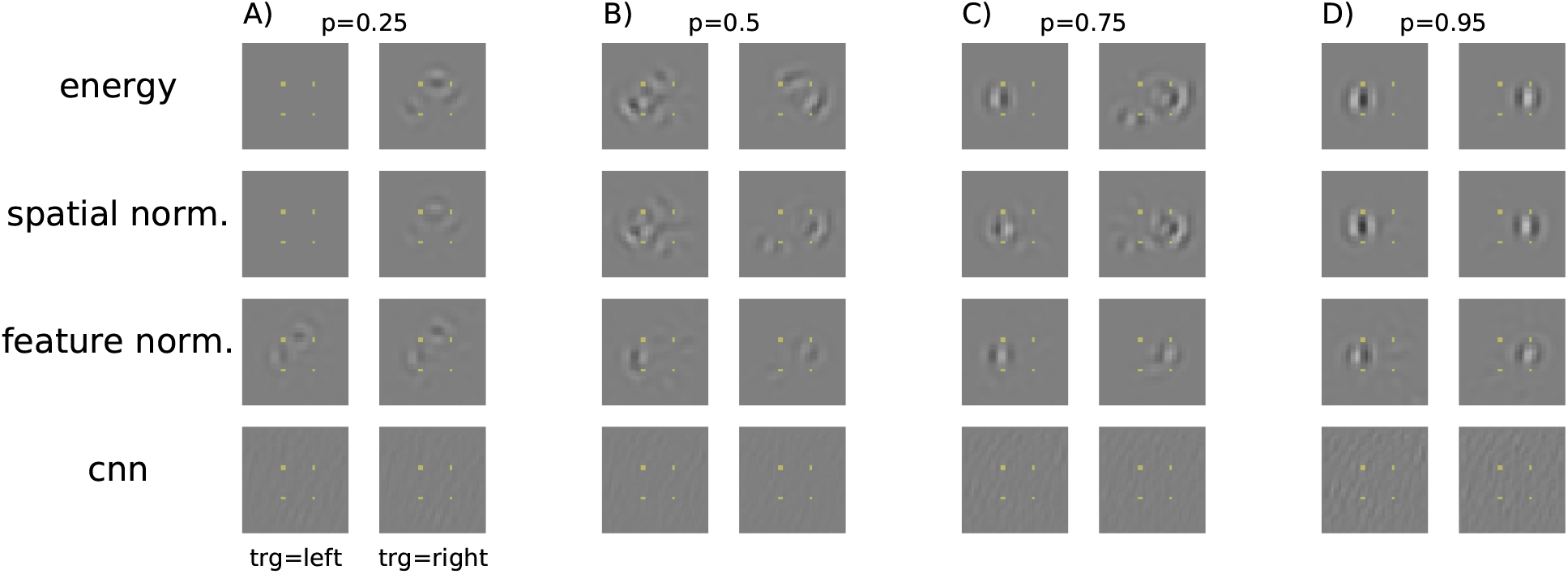
Optimized stimuli to target different performances of observer o1. (A) Stimuli targeting a performance of 25% correct responses. For reference, the target marker is shown as an overlay. Stimuli in the left column require a “left” response, stimuli in the right column require a “right” response. Different rows correspond to different models. (B) Stimuli targeting a performance of 50% correct responses. Otherwise like (A). (C) Stimuli targeting a performance of 75% correct responses. Otherwise like (A). (D) Stimuli targeting a performance of 95% correct responses. Otherwise like (A).

As the optimized stimuli in Figure 6 target higher performance levels, we find that structure in the images in the first three rows becomes more focused in the target region (i.e. left or right in Figure 6). Furthermore, images that target high performance levels (Figure 6D) have higher contrast than images that target chance performance (Figure 6A) for the models shown in the first three rows. While the first three rows of Figure 6 show stimuli for models that are—to some extent—inspired by knowledge about the early visual system, the model in the fourth row of Figure 6 is a convolutional neural network that has been optimized to correctly predict responses in the main experiment, without necessarily mimicking known properties of the visual system. Optimized stimuli for this model look quite different from stimuli for the remaining three models. In general, contrast for these images was low and did not change with the target performance level.

When observers repeated the experiment with these optimized stimuli, their performance was indeed correlated to the performances predicted by the model for the first three models (average correlation for Energy model *r* = 0.81, Spatial Normalization model *r* = 0.82, Feature Normalization model *r* = 0.89, see Figure 7A-C). However, humans tended to perform better than predicted by models based on oriented stimulus energy consistent with the idea that humans have access to additional, potentially more high-level image features than these simple models (see Figure 7A-C). On the contrary, human accuracy hardly correlated with the performance predicted by the convolutional neural network model (*r* = *−*0.18, see also Appendix “Kernels for first layer of CNN”). Even for stimuli where the neural network model predicted near perfect accuracy (p=0.95), human accuracy was still close to chance (o1: 36.8*±*3.8% human accuracy *±* s.e.m., o2: 25.0*±*3.4%, o3: 30.0*±*3.6%).

**Figure 7:**
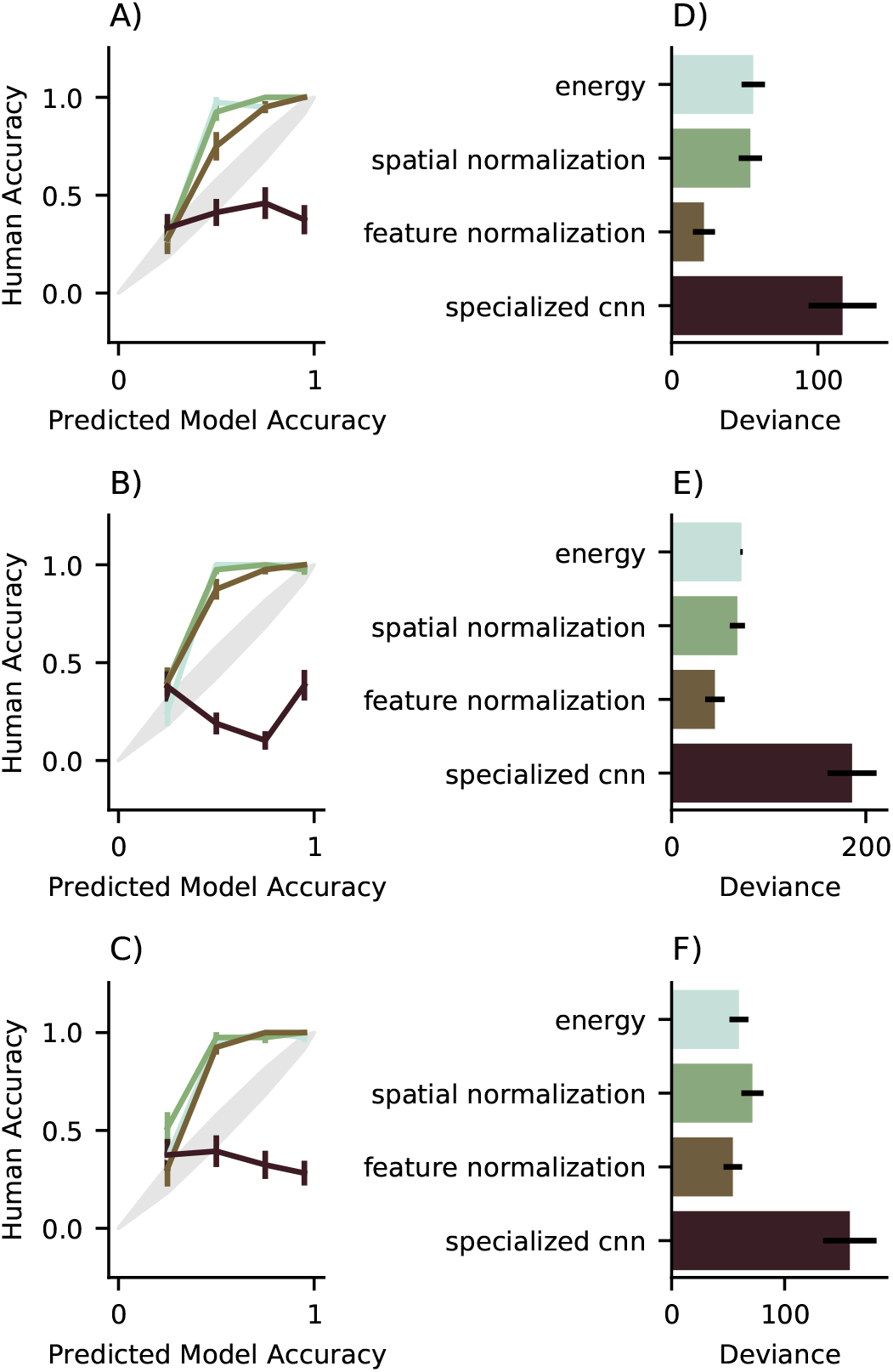
Human performance on optimized stimuli (A) Performance of observer o1 for stimuli that target different accuracies in the models. Error bars indicate standard error of mean. The diagonal gray line indicates equal performance between predicted and human (width of the line is *±*s.e.m.). (B) and (C) same as (A) for observers o2 and o3. (D) Deviance between predicted and observed performance for stimuli optimized for different models of observer o1’s trial by trial behaviour. (E) and (F) same as (D) for observers o2 and o3.

We quantified the differences between the different models by calculating the deviance between observed and predicted accuracy for each model. Deviance is a generalization of the well known sum-of-squares error to a setting with binomial responses (Dobson & Barnett, 2008). Figure 7D-F shows deviances for four different models. For observer o1 (Figure 7D), the deviance between observed and predicted accuracy was lowest for the Feature Normalization model where the normalising signal was pooled over different orientations (see Table 1) and it was much larger for any of the other models (Feature Normalization vs any other model, *|t| >* 2.8, *p <* 0.01, corrected for multiple comparisons using the method by Benjamini & Hochberg, 1995).

**Table 1:** Deviances relative to the predicted accuracies. Abbreviations are d.f.=Degrees of freedom, s.e.=standard error. Standard errors were determined using 1000 bootstrap samples.

| Observer | Model | Deviance | d.f. | s.e. |
| --- | --- | --- | --- | --- |
| o1 | Energy | 55.84 | 153 | 2.69 |
|  | Spatial Normalization | 53.87 | 155 | 2.84 |
|  | Feature Normalization | 22.13 | 160 | 2.72 |
|  | CNN | 116.92 | 150 | 2.69 |
| o2 | Energy | 71.98 | 160 | 2.70 |
|  | Spatial Normalization | 67.69 | 160 | 2.53 |
|  | Feature Normalization | 44.58 | 160 | 2.76 |
|  | CNN | 185.81 | 152 | 2.72 |
| o3 | Energy | 59.56 | 156 | 2.79 |
|  | Spatial Normalization | 71.48 | 157 | 2.59 |
|  | Feature Normalization | 54.17 | 155 | 2.72 |
|  | CNN | 157.68 | 141 | 2.89 |

For the other observers (Figure 7B,E and C,F), a similar pattern emerged: While the predicted accuracies for the CNN model were largely unrelated to the observed human accuracy, the other models generated stimuli for which the predicted accuracy increased with human accuracy. However, and similar to observer o1, observers o2 and o3 showed higher accuracy than expected from the respective models. Looking at deviances, we found a similar pattern as well (see Table 1). For both observers, the CNN model was worse than other models (o2: *D* = 185.81 *±* 2.72, *|t| >* 4.5, *p <* 10^*−*4^ comparing to other models, o3: *D* = 157.68 *±* 2.89, *|t| >* 3.5, *p <* 0.002 comparing to other models, all *p*-values corrected using the method by Benjamini & Hochberg, 1995). Furthermore, the Feature Normalization model was better or equally good than the other biologically inspired models (o2, *t*-test Feature Normalization vs Energy, *t*(318) = *−*2.96*, p <* 0.05, *t*-test Feature Normalization vs Spatial Normalization, *t*(318) = *−*1.92, *n.s.*, o3, *t*-test Feature Normalization vs Energy, *t*(309) = 0.46, *n.s.*, Feature Normalization vs Spatial Normalization, *t*(308) = 1.34, *n.s.*, all *p*-values corrected using the method by Benjamini & Hochberg, 1995).

This suggests that overall a model in which the output of the Feature Normalization model is used to derive a decision provides the best explanation for the edge localization task considered here. This model consists of energy filters with similar orientation tuning as complex cells in primary visual cortex, followed by biologically realistic gain control pooled over multiple different orientations.

### Feature weighting for the Feature Normalization model

In order to understand how features from different image locations contributed to the observers’ decisions, we visualized the readout weights of the Feature Normalization model. The readout weights map from 4 orientations at each image location to one of four different responses. We decided to visualize the readout weights for each combination of orientation and response separately (see Figure 8). Thus, in order to derive a decision signal for a “left” response, the model would match the four maps in the first row of Figure 8 and sum up the corresponding responses.

**Figure 8:**
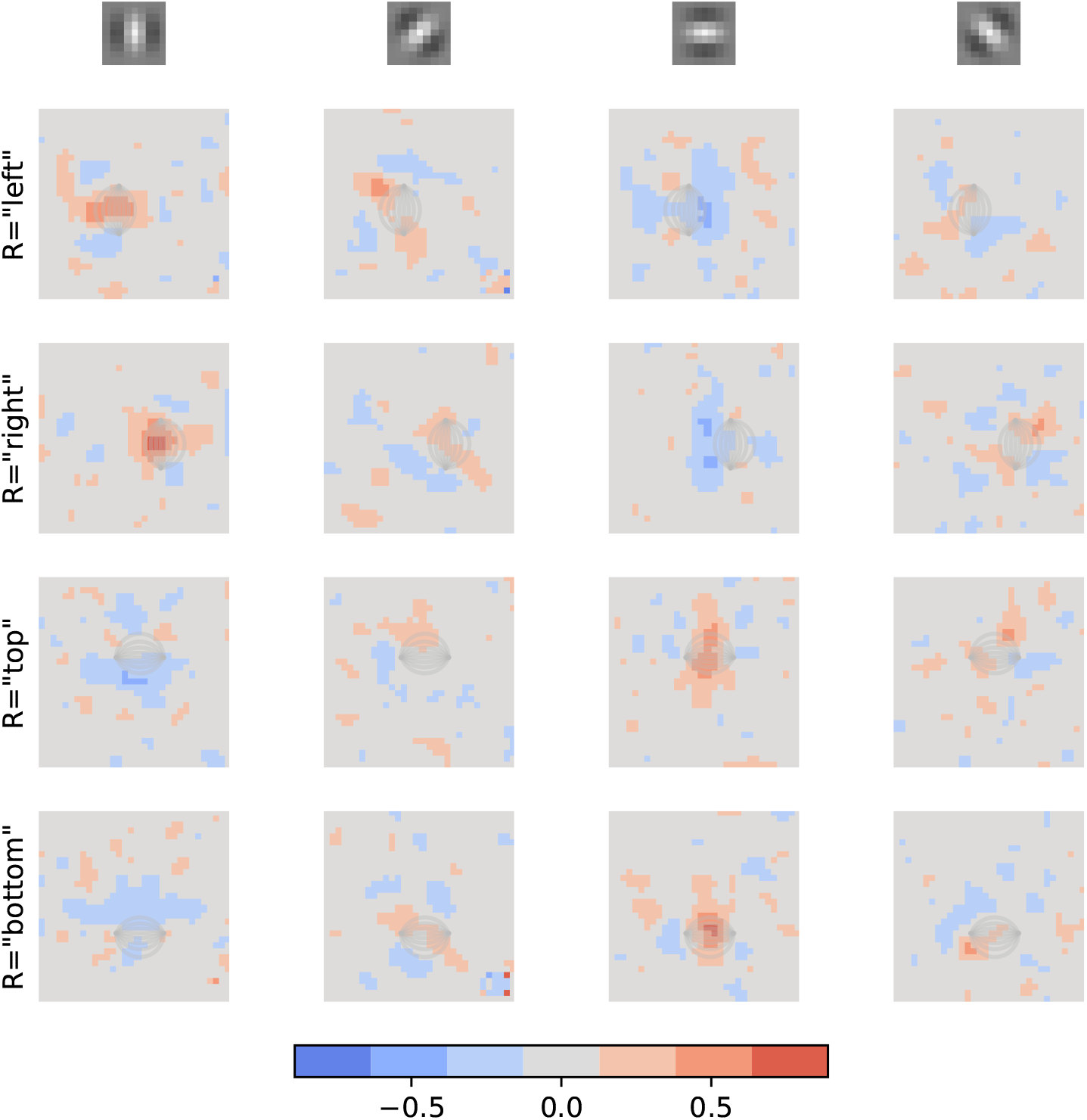
Readout weights for Feature Normalization model. Each column corresponds to one oriented energy feature, each row corresponds to one possible response. The orientation of the corresponding energy features is given by the small grating symbols above the columns. Color codes the weight with which the respective location contributed to the observer’s decision. On each panel, the arcs that would be associated with the corresponding decision are superimposed in light gray.

Figure 8 visualizes the weights for one observer (observer o2, other observers had similar patterns but they were more noisy and are shown in Appendix “Readout weights for observers o1 and o3”). Three things are worth pointing out. (i) We found that for the left and right responses vertical orientations in the area of the target signal were strongly weighted and for the top and bottom responses, horizontal orientations in the area of the target signal were strongly weighted. In the following, we will refer to this weight pattern as *ρ*_1_ (see Figure 9A). These weights correspond to signals that are aligned with a straight line interpolation between the start and end points of the respective arc. (ii) The model negatively weighted signals that corresponded to structure that was orthogonal to this straight line interpolation, but only towards the inside of the square formed by the four target markers (*ρ*_2_, see Figure 9B). (iii) We find some weight associated with oblique directions (*ρ*_3_, see Figure 9C). Specifically, locations in which curved arcs would connect to the corresponding target markers were positively weighted to arrive at a response.

**Figure 9:**
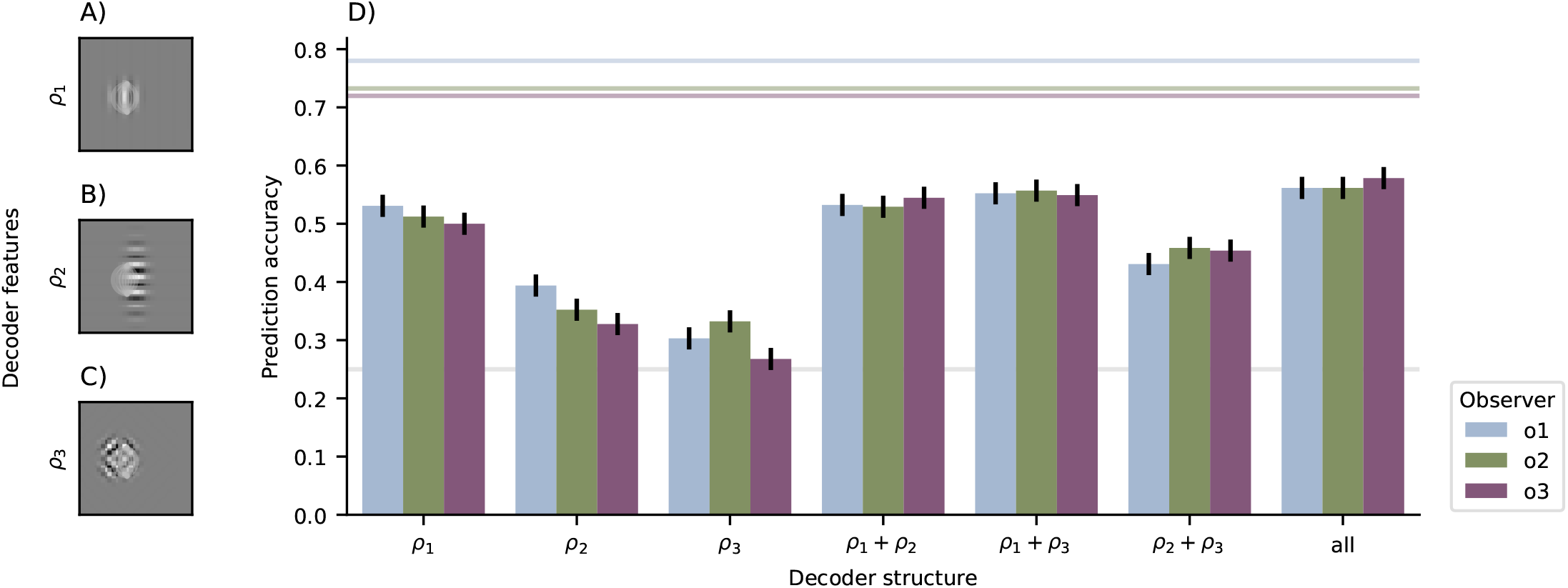
Performance of models with simplified decoder structure. (A) Schematic visualization of the decoder weights for weight pattern *ρ*_1_ for response “left”. The weight pattern was just the envelope of the pattern shown here, applied to the orientation channels visualized by the underlying grating. For reference, the superimposed lines indicate where the corresponding target was located. (B) Same as (A) for weight pattern *ρ*_2_. Note that this pattern was associated with negative weights. (C) Same as (A) for weight pattern *ρ*_3_. (D) Accuracy of prediction of human responses for the different weight patterns in isolation and combined. Similar to Figure 5, the horizontal lines indicate change performance (gray) and double pass consistency for the individual observers (colored lines).

In order to test the relevance of this visible structure in the readout weights, we created simplified models in which either of the above weight patterns were approximated by Gaussian blobs and the others were set to 0 (see Appendix “Simplified models of readout weights” for details). The predictive performance of these simplified models (and their combinations) was then evaluated on the test dataset from experiment 1 (see Figure 9). Weight pattern *ρ*_1_ predicted human responses with an accuracy of about 50% (mean*±*s.e.m.=53.1*±*1.95% for observer o1, 51.2*±*1.96% for observer o2, 50.0*±*1.96% for observer o3). Combining weight pattern *ρ*_1_ with either of the other two patterns predicted human responses with higher accuracy and combining all three components of the weight pattern predicted human performance even better (see Table 2). However, compared to the results presented in Section “Prediction performance can not discriminate between different models”, these numbers are pretty low, suggesting that the detailed pattern of feature weights really matters for a full explanation of behaviour.

**Table 2:** Performance of models with simplified decoder structure. Reported values are mean*±*s.e.m. on hold out test dataset.

| Decoder structure | Accuracy |  |  |
| --- | --- | --- | --- |
|  | o1 | o2 | o3 |
| $\rho_1$ | $0.531 \pm 0.020$ | $0.512 \pm 0.020$ | $0.500 \pm 0.020$ |
| $\rho_2$ | $0.303 \pm 0.018$ | $0.332 \pm 0.018$ | $0.268 \pm 0.017$ |
| $\rho_3$ | $0.394 \pm 0.019$ | $0.352 \pm 0.019$ | $0.328 \pm 0.018$ |
| $\rho_1 + \rho_2$ | $0.532 \pm 0.020$ | $0.529 \pm 0.020$ | $0.545 \pm 0.020$ |
| $\rho_1 + \rho_3$ | $0.552 \pm 0.020$ | $0.557 \pm 0.019$ | $0.549 \pm 0.020$ |
| $\rho_2 + \rho_3$ | $0.431 \pm 0.019$ | $0.458 \pm 0.020$ | $0.451 \pm 0.020$ |
| all | $0.562 \pm 0.019$ | $0.562 \pm 0.019$ | $0.578 \pm 0.019$ |

## Discussion

By embedding edge segments in samples from a generative adversarial network, we were able to construct edge stimuli with considerable variation in appearance. Using these stimuli, we found that human performance at detecting these embedded edge segments is consistent with multiple variations of the standard model of early vision (Schütt & Wichmann, 2017), as well as an artificial neural network model with no explicit knowledge about the visual system. We therefore constructed stimuli that, for each one of the models, would target specific levels of performance. When tested on these model specific stimuli, we found that the standard model generally performed better than the artificial neural network model.

This study relied on the CIFAR10 dataset (Krizhevsky, 2009), which consists of fairly small images from 10 different classes. It might be argued that this dataset does not appropriately cover the variability in appearance or class identity covered by larger datasets with higher resolution, such as the ImageNet dataset (Russakovsky et al., 2015). There are two perspectives on this. Firstly, one might ask to what extend the statistical regularities of the images in CIFAR10 carry over to larger images and datasets with more variability. Zoph et al. (2018) improved classification performance on the large, high resolution ImageNet dataset by optimizing hyperparameters of their network architecture on the much smaller CIFAR10 dataset. This suggests that at least part of the statistical regularities that can be learned from CIFAR10 carry over to larger and more complex image databases. Secondly, one might wonder if the statistical regularities present on CIFAR10 are consistent with the kinds of image features that normal observers use during everyday vision. The results in Appendix “Naturalism of CIFAR10 images” suggest that this is true as well. Together this supports the idea that our results would be expected to generalize to images with larger variability in appearance and class identity.

On the model specific stimuli, none of the models perfectly generalize from the naturalistic stimuli to the targeted stimuli constructed for those models. Although the biologically inspired models result in stimuli for which human accuracy increases with model accuracy, humans actually perform better for these stimuli than expected from the respective models. We believe that this implies that human observers have access to more complex interactions between signals at different locations than any of the models studied here. One class of such interactions might be effects related to surround suppression (Cavanaugh et al., 2002), where the output of a neuron with receptive field in one location is normalized by the output of neurons with neighbouring receptive fields (Carandini & Heeger, 2012; Coen-Cagli et al., 2012). Our Feature Normalization model contains a normalizing mechanism that would result to some extent in surround suppression, yet recent evidence suggests that realistic surround suppression would likely be considerably more complex (Coen-Cagli et al., 2015). Furthermore, it might be that correlations between neural responses (Kohn et al., 2016) or flexible assignment of neurons to different suppressive pools (Guerrero-Colón et al., 2008) could play a role. We believe that a detailed examination of the contributions of surround suppression to observers’ performance in our experiment is beyond the scope of this paper and decided to restrict ourselves to better understood local gain control operations.

Many studies that aim to identify features that contribute to an observers decision have used white noise (Gold et al., 2000; Abbey & Eckstein, 2002; Morgenstern & Elder, 2012; Neri, 2009, 2015; Neri & Levi, 2008, see Murray, 2011 for an overview) or noise with relatively simple correlation structure (Wilder et al., 2018). The embedding technique used here can be interpreted as “natural image noise”^2^. The convolutional neural network model fails to generalize to stimuli synthesized for this model. This could be interpreted as a failure to generalize from natural noise to less natural noise, which might seem less critical than a failure to generalize from artificial stimuli to natural conditions. However, looking at Figure 6, the images generated for the convolutional neural network generally have fairly low contrast and even for stimuli that should be recognized with high confidence, the image structures are very weak. It might be that the CNN model in our study does not learn the same features that humans use to solve the task (also see Appendix “Kernels for first layer of CNN”). Baker et al. (2018) find that convolutional neural networks trained on a large database of natural images (Russakovsky et al., 2015) use different image features than humans for object recognition. More specifically, Geirhos et al. (2019) report that such convolutional neural network models mostly rely on texture to perform classifications, while humans rely more on object shape. The CNN model here was not trained to perform large scale image recognition, but it was explicitly trained to predict human responses. However, if there was a correlation between the features used by our human observers and some subtle texture properties in the images from the main experiment, the CNN model might be biased to pick up these texture features. When constructing stimuli from the CNN model, these stimuli would only contain the subtle texture properties but not the features used by our human observers, resulting in near chance performance as observed in our control experiment.

Convolutional neural networks that have been pre-trained on the ImageNet database (Russakovsky et al., 2015) are commonly used as generic features in both computer vision (Gatys et al., 2016; Xie & Richmond, 2016, also see He et al., 2019) and biological vision (Baker et al., 2018; Khaligh-Razavi & Kriegeskorte, 2014; Geirhos et al., 2018; Tang et al., 2018). In our task, these models fall short at predicting human responses, in particular when compared to a simple energy detector (see Figure 5). While this might initially seem surprising given the success that these models had in other domains, it should be noted that the convolutional neural networks tested here appear to be quite insensitive to the precise spatial structure of an image (Brendel & Bethge, 2019). By being convolutional, these networks have to learn a set of features that applies equally across the entire image. The task used here requires localization of the target stimulus and might thus be particularly hard for convolutional neural networks. Indeed, extensions of convolutional neural networks for object localization (Redmon et al., 2016; Girshick et al., 2015) typically require additional mechanisms to use the convolutional features for object localization.

Our models all use a single frequency channel (albeit with different orientations), while it is well known that the human visual system has multiple frequency channels (Blakemore & Campbell, 1969; Ringach et al., 2002; Stromeyer & Julesz, 1972; Goris et al., 2013). In this respect, our models are clearly a simplification. However, we note that the models don’t seem to require this fine frequency resolution and are able to accurately predict human responses despite the limited frequency resolution. We find that adding orientation tuned gain control to the models tends to add to the model’s ability to predict human responses, and to improve the model’s ability to construct stimuli that target different performance levels. This suggests that gain control might have to some extent complemented the limited frequency resolution of our models.

Sebastian et al. (2017) report results on target detection in natural images. Their target stimulus was a grating patch of fixed size and orientation at the centre of a natural image patch. They elegantly “manipulated” properties of this background patch by selecting appropriate image patches from a large database of natural images. However, their target pattern was simply superimposed on the background image. Under natural viewing conditions, targets are usually not superimposed on the background, but they are part of the entire image as much as the background itself. Depending on the scene in which a target appears, that target may be partially occluded or be illuminated in different ways. All these factors will alter not only the appearance of the background, but also of the target itself (see Elder & Zucker, 1998 for a detailed discussion in the context of edge targets). Neri (2017) circumvents this problem, by creating small gaps in natural images at places where the target is inserted. This ensures that target appearance is not altered by the natural image background and that conclusions about local processing remain interpretable. However, it also means that this approach remains insensitive to local interactions between background and target, as the target is clearly different from the background. Our embedding approach allows us to constrain part of the appearance of the target while still maintaining the fact that the target is part of the background scene. In that sense, it can be seen as an intermediate approach to the ones by Sebastian et al. (2017) and by Neri (2017).

Our method of constructing model specific stimuli that target specific performance levels has some resemblance to the construction of maximum differentiation stimuli (Wang & Simoncelli, 2008). Wang & Simoncelli (2008) address the problem of comparing two competing models and they suggest constructing two classes of stimuli that each clip the accuracy of one model while maximizing accuracy of another model. Although it would be possible to generalize this procedure to comparisons between *n* models by clipping the accuracy of *n −* 1 models and maximizing the accuracy of the remaining one, this approach requires repeated evaluation of every model when constructing each one of the stimulus classes. In addition, clipping accuracy for *n −* 1 related models will result in complex constraints on the generated images that can be computationally costly to satisfy. In this study, we compared 4 different models, with three of them being closely related to each others. We take an alternative approach to Wang & Simoncelli (2008) by requiring models to construct stimuli that target a given level of human accuracy. This requires the models to also predict human responses at intermediate accuracy levels correctly while at the same time being less computationally demanding.

## Conclusion

In conclusion, we have provided evidence that the standard model of early vision, combined with a flexible linear readout mechanism, is able to generalize to fairly complex target detection tasks in naturalistic stimuli, while convolutional neural networks provide less general descriptions of visual behaviour.

Code and data for this study will be made available upon publication at doi:10.5281/zenodo.2535941.

## Appendix

### Naturalism of CIFAR10 images

To test if the CIFAR10 database captures those image features that a normal observer would use during “natural” vision, one reviewer suggested to test if a naïve observer would be able to correctly recognize grey scale images from CIFAR10. We asked one naïve, untrained observer to classify 500 images from CIFAR10. The images were randomly selected, converted to grey scale and presented at 8bit resolution on a linearized monitor (details of the stimulation setup were the same as in the main experiment). The task was set up as a spatial 2-alternative-forced-choice (2AFC): The observer saw two images from two different classes side by side along with the question “Which image is a XX”, where XX was replaced by the class of one of the two images. The observer then indicated which image they believed to belong to the target class by either pressing the left or right arrow key on a computer keyboard. There was no feedback.

On average, the observer correctly identified the target image in 97.4*±*0.7% (mean*±*s.e.m.), suggesting that classification was very easy. As shown in Table 3, the naïve observer could easily identify the target image for all target classes, with lowest performance for Dogs (94.1*±*3.2%). After the experiment, the observer reported that this was the first time, that they participated in a 2AFC task and that the task felt quite natural to them. The observer further reported typically being able to unambiguously identify the class of both images, with the exception of images from the “Frog” category.

**Table 3:** Classification accuracy of one observer on CIFAR10 images (converted to grey scale).

|  | Airplane | Automobile | Bird | Cat | Deer | Dog | Frog | Horse | Ship | Truck | Total |
| --- | --- | --- | --- | --- | --- | --- | --- | --- | --- | --- | --- |
| mean | 1.000 | 0.952 | 0.982 | 0.979 | 0.977 | 0.941 | 0.980 | 1.000 | 0.976 | 0.955 | 0.974 |
| se | 0.000 | 0.033 | 0.018 | 0.021 | 0.022 | 0.033 | 0.020 | 0.000 | 0.024 | 0.025 | 0.007 |

Using a 2AFC task, allowed us to measure the amount of information available to the naïve observer without taking into account a possibly suboptimal decision criterion (Green & Swets, 1966). Although the observer reported that the task felt quite natural, it is likely that they would perform worse in a more natural setting. However, the extremely high performance in this task and the subjective report of the naïve observer are consistent with the idea that the CIFAR10 dataset indeed captures information that a normal observer would use during natural vision.

### Validation of embedded arc stimuli as “natural”

Embedding arc segments clearly changes an image’s statistics *locally*; otherwise the visual system wouldn’t be able to recognize that arc segment. In order to verify that the embedded arc stimuli only affected the image’s statistics locally, but left other parts of the image untouched, we analyzed quadrants of images (see Figure 10A). A large number of statistical signatures of natural images have been described. These include a characteristic scaling law of the images’ power spectrum (Field, 1987) or second order correlations between local wavelet statistics (Wainwright & Simoncelli, 2000; Schwartz & Simoncelli, 2001). Although these image properties are well known, they are only moderately meaningful on the 16*×*16 image quadrants used here. We therefore decided to adopt a different approach: We asked if a 4-layer version of the VGG network (Simonyan & Zisserman, 2015) could tell apart quadrants from natural images, from unconditioned GAN samples and from images that had an arc segment embedded in another part, as well as images from a model that matches an image’s second order statistics (Portilla & Simoncelli, 2000, we used their code in the default configuration, but with 3 instead of 4 levels for the pyramid decomposition, due to the small image size) and images that matched the images power spectrum. Specifically, the VGG network had four convolutional layers with max-pooling after every second layer, followed by two fully connnected readout layers (in the notation by Simonyan & Zisserman (2015) the network’s architecture was conv3_64, conv3_64, maxpool, conv3_128, conv3_128, maxpool, FC_64, FC 3, softmax). We selected 1824 image quadrants per category and randomly selected 400 of them as validation and test sets respectively. Thus, the remaining training set consisted of 8320 image quadrants. We trained the network for up 1000 epochs with a learning rate of 0.002 and a batch size of 128. We used early stopping to select the network with lowest cross entropy on the validation dataset (after 302 epochs). We report results for this network on the remaining test set of 400 image patches.

**Figure 10:**
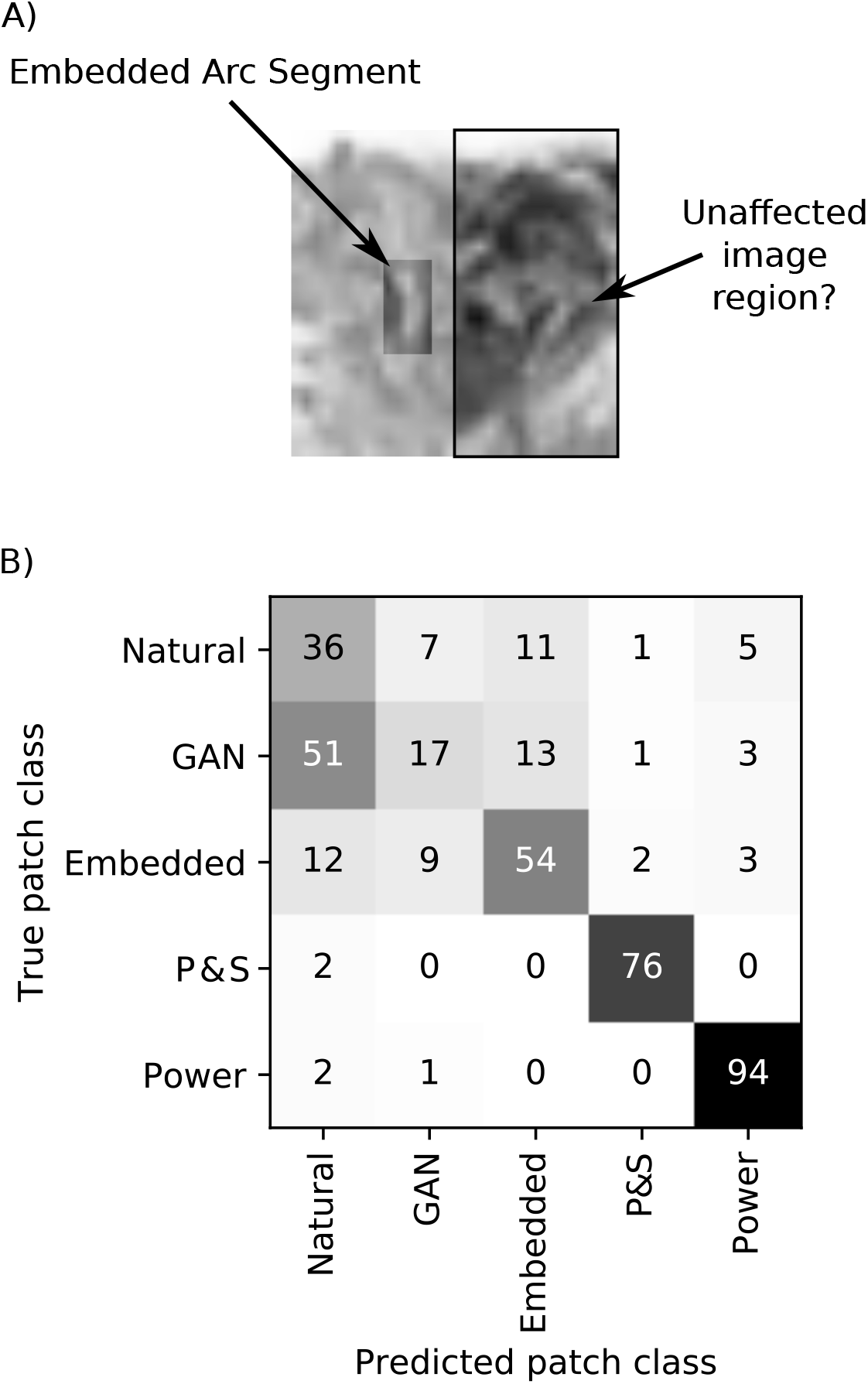
Do images with embedded arc segments have similar statistical properties as natural images? (A) The left side of the image contains an embedded arc segment, affecting the image’s statistics. To understand if the effect of this manipulation also affected the rest of the image, we analyzed a quadrant from the supposedly unaffected side of the image (right side in this example). (B) Confusion matrix of the patch classification network. While natural images and GAN samples appear mostly natural to the network, samples from the texture model by Portilla & Simoncelli (Portilla & Simoncelli, 2000, P&S) and images with matched power spectrum (Power) can be clearly told apart from natural images. Results for embedded arc images (Embedded) are somewhat in between.

Overall, the network achieved a prediction accuracy of 69.2*±*2.3% (mean *±* s.e.m.), significantly better than chance (25%). The full confusion matrix for the 5 classes reveals additional information (Figure 10B). While the network often confused GAN samples with natural images, it could easily tell apart images with the power spectrum matched to natural images from real natural images. The same was true of samples from the texture model by Portilla & Simoncelli (2000). Images with embedded arc images were somewhere in between; the network was clearly better at telling them apart from natural images than it was for the unconditioned GAN samples, but it confused embedded arc images more often with real natural images than samples from the Portilla & Simoncelli (2000)-model or images with matched power spectrum. We therefore conclude that the images used in experiment 1 are reasonably well matched to the statistics of natural images and—more importantly—match those statistics better than alternative image models would.

### Details of model architectures

#### Simple cell filter bank

The model consisted of a bank of linear, oriented filters. Filters were polar separable in the Fourier domain, such that the frequency response could be written as

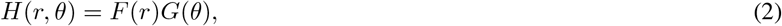

where 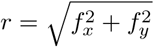 and *θ* = arctan2(*f*_*x*_,*f*_*y*_) are the (absolute) frequency and orientation. We fixed

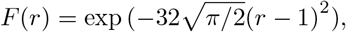

to cover approximately 1.6 octaves (Zhaoping, 2014) and selected four different orientation response functions *G*_*n*_, *n* = 0, 1, 2, 3 of the form (circularly wrapped)

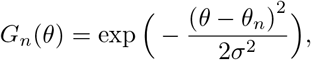

where *θ*_*n*_ = *nπ*/4 and

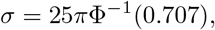

to achieve an orientation bandwidth of 20–30 degrees (Ringach et al., 2002). Here, Φ: ℝ *→* (0, 1) is the cumulative distribution function of the standard normal distribution. The filter *H* was subsequently converted to the spatial domain and only the real (even) part was retained. The resulting filter *h*_*n*_ was pruned to only contain the central 7*×*7 coefficients.

All four filters were applied to each image with zero padding at the borders, resulting in a 4 *×* 32 *×* 32 array. This array was treated as one long 4096 element vector ***s*** and was submitted to a 4-class logistic regression.

#### Complex cell filter bank

The complex cell filterbank model used the same filterbank from equation (2). However, when converting *H* to the spatial domain the real (even) and imaginary (odd) parts were retained. These were pruned in the same way as for the simple cell filter bank and they were then both applied to the input image (with zero padding). This ultimately resulted in two 4096-dimensional feature vectors ***ψ***_even_ = ***s*** and ***ψ***_odd_. From these two we constructed a local, oriented energy signal (e.g. Morrone & Burr, 1988; Adelson & Bergen, 1985) as

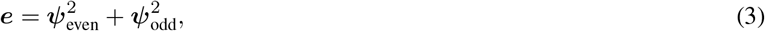

to be submitted to the subsequent 4-class logistic regression.

#### Simple cell filter bank with local gain control by simple cell outputs

This model used the same filterbank model as in equation (2) and calculated the same feature vector ***s*** as above. In addition, a normalization signal ***m*** was computed by convolving the oriented real parts of the filter outputs ***s*** with filters rotated by 90 degree. In other words, if ***s***_0_ denotes the oriented energy at orientation *θ*_0_ = 0 and *h*_2_ denotes the filter with orientation *θ*_2_ = *π/*2, then

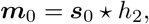

with *\** denoting 2d-convolution, thus the resulting vector ***m***_*n*_ of normalization strengths at orientation *n* had the form

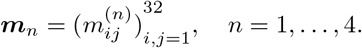

We then calculated a normalized vector of the form

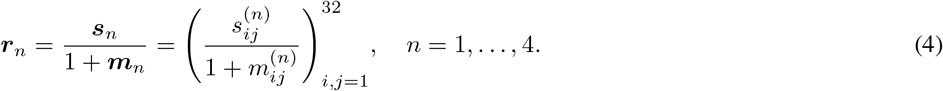

The resulting 4096-dimensional feature vector was submitted to 4-class logistic regression.

#### Simple cell filter bank with local gain control by complex cell outputs

This model was very similar to the model in the section above about gain normalized simple cell filters except that the normalization was calculated from the oriented energy signal rather than from the simple cell outputs. Thus vectors of the form

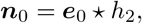

were used as a normalization signal and we calculated the vector

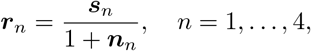

similar to equation (4). The vectors ***r*** were concatenated into one 4096-dimensional feature vector and submitted to 4-class logistic regression.

#### Complex cell filter bank with local gain control by complex cell outputs

This model was almost the same as the model in the previous paragraph except that the vectors

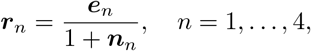

were submitted to 4-class logistic regression.

#### Complex cell filter bank with orientation tuned gain control by complex cell outputs

We often think of the gain control signal as being tuned both in location and orientation, where similar orientations in similar locations contribute most to the normalization pool. We therefore used a model in which the gain control signal was pooled by a space-orientation separable filter of the form

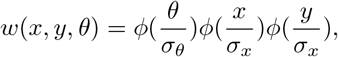

with orientation bandwidth *σ*_*θ*_ = *π*/2 and spatial bandwidth *σ*_*x*_ = 1 pixel (Schütt & Wichmann, 2017). Here, *φ*: ℝ → ℝ was the density function of the standard normal distribution. We then used

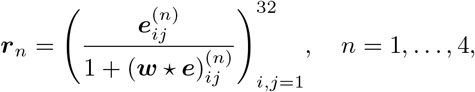

in our 4-class logistic regression. Note, that here a 3d-convolution is taken between the 4 *×* 32 *×* 32 orientation by space by space tensors ***w*** and ***e***. That 3d-convolution treated the orientation dimension as circular and the two spatial dimensions were padded by zeros.

#### Deep convolutional neural networks

We studied two types of convolutional neural networks. Firstly, we used pretrained versions of VGG (Simonyan & Zisserman, 2015) and ResNet (He et al., 2015) in which we replaced the final fully connected layers by a single linear decoding layer. The networks were downloaded through torchvision and the final decoding layer was trained to predict the observers trial by trial responses using stochastic gradient descent with a learning rate of 0.001 and a batch size of 16 trials. In order to make the pretrained networks compatible with the small images used in this study, we first scaled up each image by a factor of two to a size of 64 *×* 64 pixels using bilinear interpolation.

Secondly, we specifically trained a full convolutional neural network to predict human responses with approximately the same accuracy found to be double pass consistency in our observers. This deep convolutional neural network constructed non-linear features in 5 convolutional layers. Networks with fewer layers had lower performance. Each convolutional layer had 4 output channels and a learned kernel of size 3 *×* 3. The convolution operation was followed by rectification (Glorot et al., 2011) and batch normalization (Ioffe & Szegedy, 2015). The resulting non-linear features were then cast into one long vector of 4096 values and were linearly read out using logistic regression as in the other models. In total, the deep convolutional neural network had 17 060 parameters. Thus, fitting this network was really dependent on using effective regularization techniques. Batch normalization is known to have a regularizing effect (Ioffe & Szegedy, 2015) and we further used early stopping based on prediction error on a separate validation dataset (see for example Goodfellow et al., 2016, section 7.8). Although early stopping was used for the other models as well, it was particularly important for the deep convolutional neural network model.

### Kernels for first layer of CNN

We trained one convolutional neural network model specifically to predict human responses. This model successfully predicted human trial-by-trial responses in the main task, but generated targeted stimuli that did not drive human behaviour in a meaningful way. In order to gain a better understanding of what this model had learned about the images, we visualized the kernels of that model’s first layer in Figure 11. These kernels appear mostly random. Thus, although this model successfully predicted neural responses on held out test data, it appears that this model was not able to pick up consistent and reasonably smooth patterns from the input.

**Figure 11:**
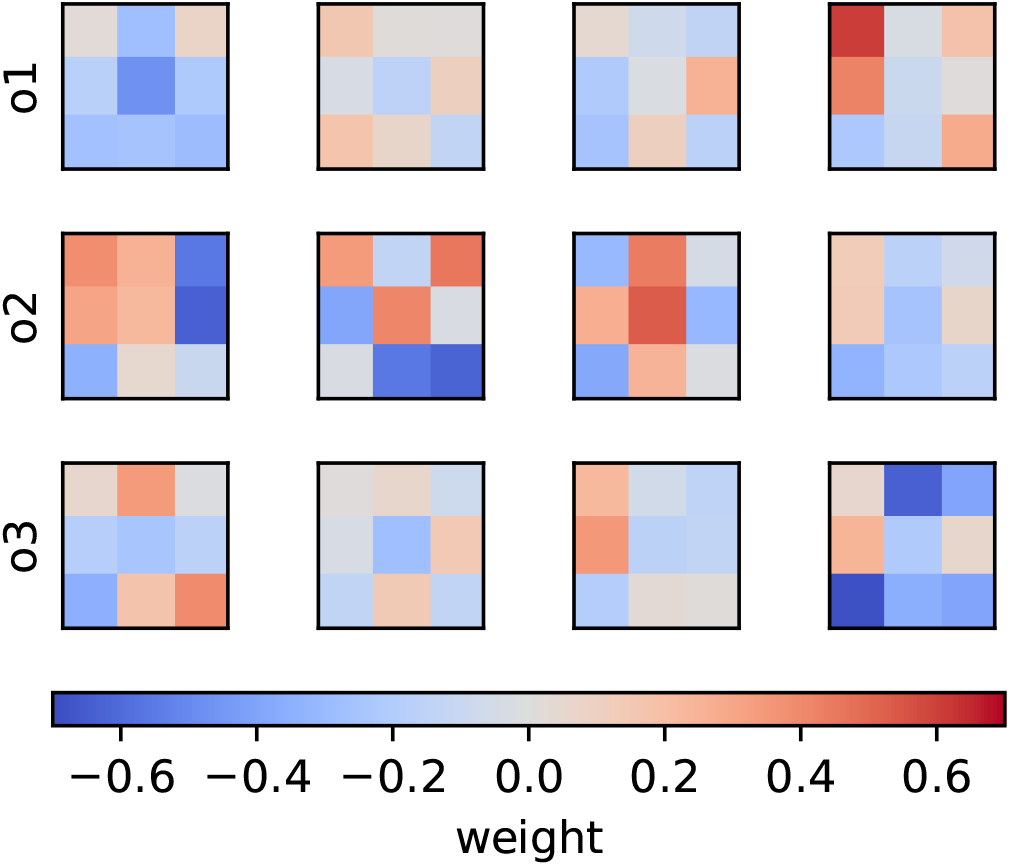
Kernels in the first layer of the neural network trained to predict human responses. Each row corresponds to one observer, each column to one of the four different kernels. Kernel weight is coded by color.

### Readout weights for observers o1 and o3

Figure 12 shows readout weights for observers o1 and o3. For both observers, the pattern of readout weights was more noisy than for observer o2. However, we found qualitatively similar structure.

**Figure 12:**
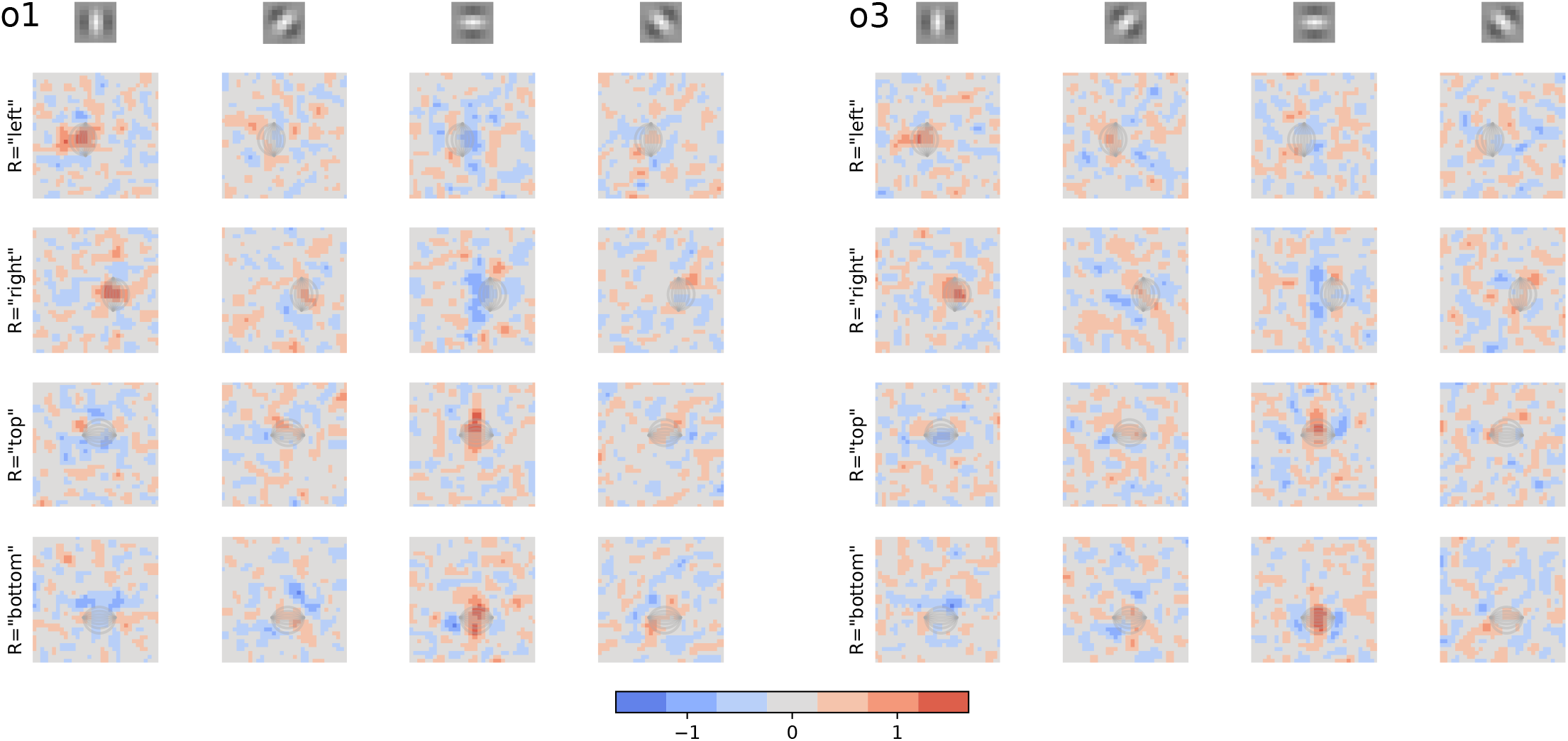
Readout weights for Feature Normalization model for observer o1 (left) and observer o3 (right). Details of the subfigures are the same as in Figure 8.

### Simplified models of readout weights

We analyzed three different decoder features and their combinations to understand the readout process for the Feature Normalization model. Specifically, those were *ρ*_1_ horizontal/vertical structure aligned with the straight line version of the target, *ρ*_2_ horizontal/vertical structure orthogonal to the straight line version of the target, *ρ*_3_ oblique structure along the sides of the arcs. To describe these features, we can think of the decoding weights as a template tensor 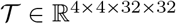, such that 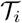 denotes the templates for response *i* (i.e. one row in Figure 8). Each of these templates has 4 orientation bands, such that 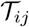 denotes the template for response *i* for features from the orientation band *j*. Note that this template is sensitive to orientations of approximately *πj*/4. Components within orientation bands are approximated by Gaussian blobs of the form *u*: ℝ^2^ × ℝ → ℝ, where

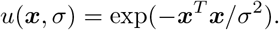

In all cases, the exact readout weights were determined using numerical optimizaton as outlined below, however the results remained qualitatively similar if the readout weights were selected using the starting values of the parameters (see below).

#### Horizontal/vertical structure aligned with the straight line version of the target

For decoding features *ρ*_1_, the template tensor 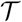 was zero except

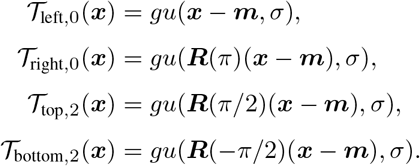

Here, we denote by 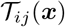 the value of the template for response *i* at orientation band *j* and location ***x***. Furthermore, we used the rotation matrix

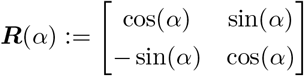

The parameters ***m***, *g*, and *σ* were estimated by minimizing the squared difference between 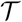 and the observed decoding weights. The minimization was done using simplex search (Nelder & Mead, 1965) starting at 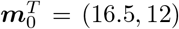, *σ*_0_ = 3, and *g*_0_ equal to the maximum of the original decoding weights.

#### Horizontal/vertical structure orthogonal to the straight line version of the target

For decoding features *ρ*_2_, the template tensor 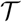 was zero except

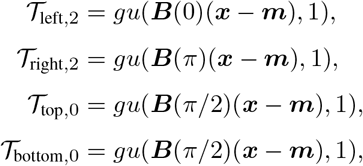

with the rotation matrix

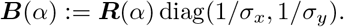

The parameters ***m***, *g*, *σ*_*x*_, and *σ*_*y*_ were estimated in the same way as in the previous section, but starting from 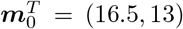, *σ*_*x*,0_ = 3, *σ*_*y*,0_ = 5 and *g*_0_ set to the minimum of the original decoding weights.

#### Oblique structure along the sides of the arcs

For decoding features *ρ*_3_, the template tensor 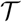 was

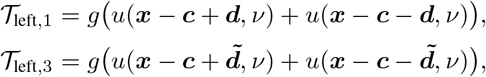

with analogous expressions for other orientations and responses. Here, the free parameters are ***c***, ***d***, *g*, and *ν* and for ***d*** = (*d*_1_, *d*_2_)^*T*^ we defined 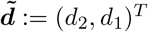. Optimization in this case started with ***c***_0_ = (16.5, 12)^*T*^, ***d***_0_ = (4, 3)^*T*^, *ν*_0_ = 2, and *g*_0_ was set to the maximum of the observed oblique feature maps.

## Acknowledgments

The author would like to thank Richard F. Murray for generously lending out several pieces of lab equipment and for feedback on an earlier version of this manuscript. This work was supported by the York University Bridging Fund and the NSERC discovery program. This research was undertaken as part of the Vision: Science to Applications program, thanks in part to funding from the Canada First Research Excellence Fund.

## Footnotes

1 In other words, the energy model was better in all 2000 permutations of the data.

2 Note however, that in this case the noise also influences the appearance of the target, and is therefore not additive as assumed in previous studies.

